# Leaving Science: Attrition of Biologists in 38 OECD Countries

**DOI:** 10.1101/2024.12.06.627162

**Authors:** Marek Kwiek, Lukasz Szymula

## Abstract

This study examines biologists leaving science in 38 OECD countries in the past two decades. We use publication metadata from a global bibliometric database (raw Scopus data at the micro-level of individual scientists). In a cohort-based and longitudinal fashion, we follow individual men and women scientists over time, from their first to their last publication (N=86,178). We examine four academic disciplines: AGRI (agricultural and biological sciences), BIO (biochemistry, genetics, and molecular biology), IMMU (immunology and microbiology), and NEURO (neuroscience). We apply survival analysis, conceptualizing scientific life as a sequence of scholarly publishing events. Our Kaplan–Meier survival analysis shows how women disappear from science: in BIO, about 60% are still in science after 5 years, 40% after 10 years, and only 20% by the end of the period examined (i.e., after 19 years). The percentages are substantially higher for men: approximately 70%, 50%, and 30%, respectively. Kaplan–Meier estimations indicate that women in the largest discipline (BIO) are 23.26% more likely to leave science after 10 years and 39.74% more likely to leave science at the end of the study period. Gender difference in attrition, slightly visible after 5 years, increase consistently in later career stages. The probability of surviving for women after 15 years varies considerably, from 47.8% in AGRI to 27.6% in IMMU; for men, the probability is about a fifth higher. Our data show that with the passage of time, women disappear from science in ever-larger proportions compared to men. Gender differences in attrition in the four disciplines have been and continue to be high, but comparing the 2000 and 2010 cohorts, have slightly decreased over time.

## Introduction

In this study, we examine biologists in 38 OECD countries leaving science in the past two decades. Our conceptual approach is straightforward: starting an academic career is making one’s first contribution to an academic journal, and leaving science is publishing one’s final research paper. We use publication metadata from a global bibliometric database (raw, structured, and curated Scopus data at the micro-level of individual scientists). We follow individual men and women scientists over time, from their first publication to their last.

Methodologically, our approach is cohort-based (Glenn, 2005) and longitudinal (Singer & Willett, 2003). We track two cohorts of scientists for up to 22 years. We examine the academic publishing careers of 34,970 biologists who started publishing in 2000 (termed the 2000 cohort); and to see the changes from a temporal perspective, we examine the publishing careers of 51,208 biologists who started publishing in 2010 (termed the 2010 cohort).

Our focus is on the four academic disciplines closely related to biology: AGRI, agricultural and biological sciences; BIO, biochemistry, genetics, and molecular biology; IMMU, immunology and microbiology; and NEURO, neuroscience. The largest discipline studied is BIO, with approximately 23,000 scientists in the older cohort and 32,000 scientists in the younger cohort. In total, we analyze the publishing histories of 86,178 scientists from 38 OECD countries, tracking the entirety of their academic careers until 2022.

We quantify the phenomenon commonly referred to in the literature as “leaving science” (Rosser, 2004; Xu, 2008; Preston, 2004; Geuna & Shibayama, 2015; Kaminski & Geisler, 2012; White-Lewis et al., 2023). Our original, larger study was focused on 16 STEMM disciplines (science, technology, engineering, mathematics, and medicine) and 11 non-overlapping cohorts of scientists (N=2,127,803; Kwiek & Szymula, 2024). The present study is restricted to the four related disciplines and two cohorts of scientists.

We use micro-level bibliometric metadata regarded as the digital traces left by (publishing) academic scientists over their careers. Consequently, academic non-publishers are not included in our research – as they leave behind no metadata of a longitudinal nature to examine. In fact, “academic career” as defined by entering and leaving science here means “publishing academic career.” However, publishing research articles in STEMM disciplines, following the long tradition of academic career studies, is equivalent to being an active researcher in the field (Hermanowicz, 2012; Leisyte & Dee, 2012).

With digital traces left behind and indexed in global publication datasets, individual scientists can be studied according to their academic age, seniority, institution type, collaboration and publishing patterns, mobility, as well as discipline and gender (Kashyap et al., 2023; Kwiek & Roszka, 2024), with the latter two being the focus of this study. We use a unique opportunity to track individuals over their academic careers at a level of detail and at a scale previously unattainable (Wang & Barabási, 2021). For large-scale multi-country examinations of the science profession, only bibliometric datasets are possible sources of reliable data, despite their limitations – which are widely discussed in the scientometric literature.

Our global and longitudinal approach to academic careers represents a more general turn toward structured big data in social science research (Liu et al., 2023), with repurposing of new data (Salganik, 2018). In particular, we make use of remarkable progress in defining the gender of scientists with massive gender-detection tools (Karimi et al., 2016; Sebo, 2021). Global academic career research can offer insight into the professional trajectories of men and women scientists and identify disciplinary and gender variations.

In this study, our interests go beyond biologists in any single national science system or in any given year, as in survey- and interview-based research. Our methodological approach is longitudinal (rather than cross-sectional) and global (rather than national). We view the whole traditional monolith of STEMM science as divided into several segments with vastly different representations of women, with these changing at different speeds in the past 30 years (Kwiek & Szymula, 2023).

Our original research about 16 STEMM disciplines (Kwiek & Szymula, 2024), from which the data on biologists are drawn, has been discussed in 25 countries and in 15 languages, as well as featured in *Nature* (Naddaf, 2024). We assume that the response to this research, in both popular and professional venues, is related to its scale, scope, and longitudinal dataset, which was used for new purposes – examining the academic profession at a large scale, beyond national borders, and by gender.

In the present study, we turned a large, raw, bibliometric dataset (Scopus) into a large dataset about academic careers in biology-related disciplines. The study is longitudinal in the strict sense of the term: The two cohorts are tracked over time for up to two decades (2000–2022) on a yearly basis. Scopus is particularly suitable for global analysis at the micro-level of individual scientists because it is structured around individual Scopus Author IDs, apart from its focus on publications and their metadata (Baas et al., 2020). In terms of data flow, our sample consists of all scientists starting to publish in 2000 or 2010, publishing in one of four disciplines (AGRI, BIO, IMMU, and NEURO), having at least two research articles in their lifetime publishing portfolios, and having clearly defined affiliations in an OECD country and a clearly defined gender (binary: male or female).

## Previous studies

Here, our focus is on persistent gender differences in biologist attrition (leaving science). Research conducted over the past three decades has shown that attrition is generally greater for women than for men (Preston, 2004; Kaminski & Geisler, 2012); women are reported to leave science earlier and much more frequently than men. Gender differences in attrition have been explained by several hypotheses and discussed using useful metaphors such as “leaky pipeline” and “chilly climate” (Blickenstaff, 2005; Goulden et al., 2011; Shaw & Stanton, 2012). In general terms, a chilly climate in STEM disciplines means a hostile or unwelcoming work environment for women which can discourage them both from entering and pursuing academic careers (Cornelius et al., 1988).

While the “pipeline” perspective is focused on individuals, the “pathways” perspective is focused on the organizational structures in which individual scientists are set (Fox & Kline, 2016). The critical point in the pathways perspective is that the social structures of the science enterprise are changeable. While the focus in the pipeline literature is on individual women in science (and on how to decrease their leakage from the scientific pipeline), the focus of the pathways literature is on the structural conditions of science in which women are embedded (Fox, 2020). A somehow passive flow of women between academic career stages is assumed in the first metaphor; in the second, in contrast, the focus is on possible actions to be taken to make academic workplaces more equitable (Xie & Shauman, 2003).

The pipeline metaphor dismisses signals about who naturally belongs to science and who is naturally excluded from it – who is the stereotypical ideal scientist (in male-dominated disciplines such as computing). It ignores gendered obstacles and disregards “persistent exclusionary messages” directed at women (Branch & Alegria, 2016). The pipeline conceptual perspective leads to the conclusion that the underrepresentation of women in science is attributable to women’s relatively low supply and their relatively higher rates of attrition from the science pipeline. As a result, the practical answer for this perspective is to block leakage (Xie & Shauman, 2003) and to increase the supply of women (as criticized in Fox & Kline, 2016).

The “pathways” metaphor, in contrast, emphasizes progression in science that is neither direct nor simple and focuses on the organizational culture in which scientists are embedded (Branch, 2016). What matters for progression in science is more than individual traits. It is the organizational culture, which involves shared values, beliefs, and behaviors; a dominating culture defines what is more (and what is less) valued in organizations and promotes standard “ways of doing business.” Ways of doing business also tend to influence how women collaborate internationally (Kwiek & Roszka, 2021a) and how they form research teams (Kwiek & Roszka, 2021b).

The conceptual frameworks employed in attrition studies include various “push” and “pull” factors. Explanations of quitting science include problems with maintaining a healthy work– life balance, low job security, low salary, colleague and workload concerns, as well as various types of discrimination in the workplace and hostile workplace climates in academic departments.

## Data and methods

We analyzed the publishing careers of 34,970 biologists who started academic publishing in 2000 and 51,208 biologists who started academic publishing in 2010. We examined four biology-related disciplines: AGRI, BIO, IMMU, and NEURO. In all, we analyzed the publishing histories of 86,178 biologists from 38 OECD countries, tracking the entirety of their publishing careers until 2022. Our data come from Scopus, a global publishing and citation database, and access to the raw Scopus dataset was granted through the International Center for the Study of Research (ICSR) Lab platform based on a multiyear collaboration agreement.

Leaving science was conceptualized as no longer publishing in academic journals. We used a “survival analysis” of the publishing behavior of individuals over the years – until the event of finally not publishing occurs. Using the bibliometric dataset, we were able to locate in time the event of not publishing and to analyze gender and disciplinary differences in attrition (and retention) for different cohorts of scientists, or for scientists starting their academic careers at different points in time.

We conceptualized scientific life as a sequence of scholarly publishing events. Scientists do research – and publish research findings. Leaving science (or quitting publishing) was therefore conceptualized as an event located in the academic biographies of scientists, and survival analysis was applied. In survival analysis, questions related to the timing of, and the time leading up to, the occurrence of an event are explored.

In our case, some scientists did not experience the event in 2022 or earlier. They continued publishing. These were termed right-censored observations about which we had only partial information (for instance, the event may have occurred in 2023 or 2024). Therefore, to classify an author as leaving science, the final publication had to be dated 2018 (and marked as leaving science in 2019) or earlier. We used only uncensored cases, which represent observations for which we have both the start and end year of being in science.

For the four disciplines, for each subsequent year, we had the initial number of scientists entering the time interval (which consisted of scientists who left science during the interval and those who stayed in science in the following interval – in our case, the following year). The probability of survival until a given time interval was calculated by multiplying all probabilities of survival across all time intervals preceding the time (Mills, 2011).

Survival analysis is presented in tables, separately for each discipline, and in graphs that highlight disciplinary differences in attrition and retention over time and by gender. We tested the power of structured, reliable, and curated big data of the bibliometric type to examine milestones in academic careers (as in Kwiek & Szymula, 2023 on young male and female scientists in 38 OECD countries).

From a disciplinary perspective, the major dividing line is between male-dominated STEMM disciplines, with participation of women scientists usually lower than 20% and very slowly increasing over time, and the STEMM disciplines in which gender balance has been achieved. We assume that gender balance in science means the participation of women at the level of 40%–60% (Kwiek and Roszka 2022).

In the four disciplines, for the two cohorts examined, gender balance has been achieved. For instance, in the largest discipline of BIO, 47.83% of the 2000 cohort and 52.75% of the 2010 cohort were women (Table 1). Our previous research showed that in gender-balanced disciplines, the attrition of women was higher than that of men. For the heavily male-dominated disciplines of MATH (mathematics), COMP (computing), ENG (engineering), and PHYS (physics and astronomy), in contrast, attrition rates for men and women were nearly equal across 38 OECD countries, with no statistically significant differences.

**Table 1:** The sample, four disciplines, two cohorts of scientists (2000 and 2010) by gender.

|  |  | Men |  | Women |  | Total |
| --- | --- | --- | --- | --- | --- | --- |
|  |  | N | % | N | % | N |
| Panel 1: the 2000 cohort | AGRI | 4,962 | 62.26 | 3,008 | 37.74 | 7,970 |
|  | BIO | 11,839 | 52.17 | 10,853 | 47.83 | 22,692 |
|  | IMMU | 805 | 45.35 | 970 | 54.65 | 1,775 |
|  | NEURO | 1,353 | 53.41 | 1,180 | 46.59 | 2,533 |
|  | Total | 18,959 | 54.22 | 16,011 | 45.78 | 34,970 |
| Panel 2: the 2010 cohort | AGRI | 6,755 | 52.81 | 6,037 | 47.19 | 12,792 |
|  | BIO | 14,905 | 47.25 | 16,637 | 52.75 | 31,542 |
|  | IMMU | 939 | 39.77 | 1,422 | 60.23 | 2,361 |
|  | NEURO | 2,153 | 47.71 | 2,360 | 52.29 | 4,513 |
|  | Total | 24,752 | 48.34 | 26,456 | 51.66 | 51,208 |

Leaving science is a traditional scholarly topic, but it has only been explored through case study research (based on surveys and interviews) or through multi-year U.S. studies of post-secondary faculty leading to single-nation results. Concepts such as “faculty departure intentions,” “intentions to leave,” and “faculty turnover” have been studied for several decades (Zhou & Volkwein, 2004; Rosser, 2004; Ehrenberg et al., 1991; Smart, 1990). Most attrition studies have focused on single academic institutions, and their geographical scope was restricted to the United States (exceptions include Milojević et al. 2018, who studied several disciplines by using data from major academic journals).

Membership in the four disciplines was determined by computing all cited references in all publications published during whole academic careers. All cited references for each individual scientist were linked to academic journals (using the ASJC journal classification list used in Scopus), and the most often occurring value (or the modal value) was selected. When two modal values occurred in computations, the individual scientist was removed from further analysis.

Consequently, we used a sample of scientists with clearly determined characteristics: they were all research-active in the four disciplines and they had a clearly determined country affiliation and an unambiguously determined gender. They were also assigned a year of exit from academic publishing. Details regarding determination of the various individual-level characteristics of the larger sample from which the subsample of biologists was derived were discussed in our original report and its appendices (Kwiek & Szymula, 2024). What is especially important is the validity of gender determination. We inferred gender from bibliometric metadata by using Namsor, one of the best gender-detection tools currently available (Namsor, 2024).

## Results

A classic statistical technique for survival analysis is the Kaplan–Meier estimate of survival (Mills, 2011), with a plot of the Kaplan–Meier estimator shown as a series of characteristically declining horizontal steps of various heights. Figure 1 shows gender differences in attrition (and retention) for the 2000 cohort of scientists separately for each discipline through Kaplan– Meier survival curves. Steps lead down over time, one step for one year, with probabilities of staying in science shown on the y-axis and the number of years spent publishing since 2000 on the x-axis. Small crosses denote right-censored observations for the three most recent years studied (i.e., 2020–2022), which were not used in computations.

**Figure 1.**
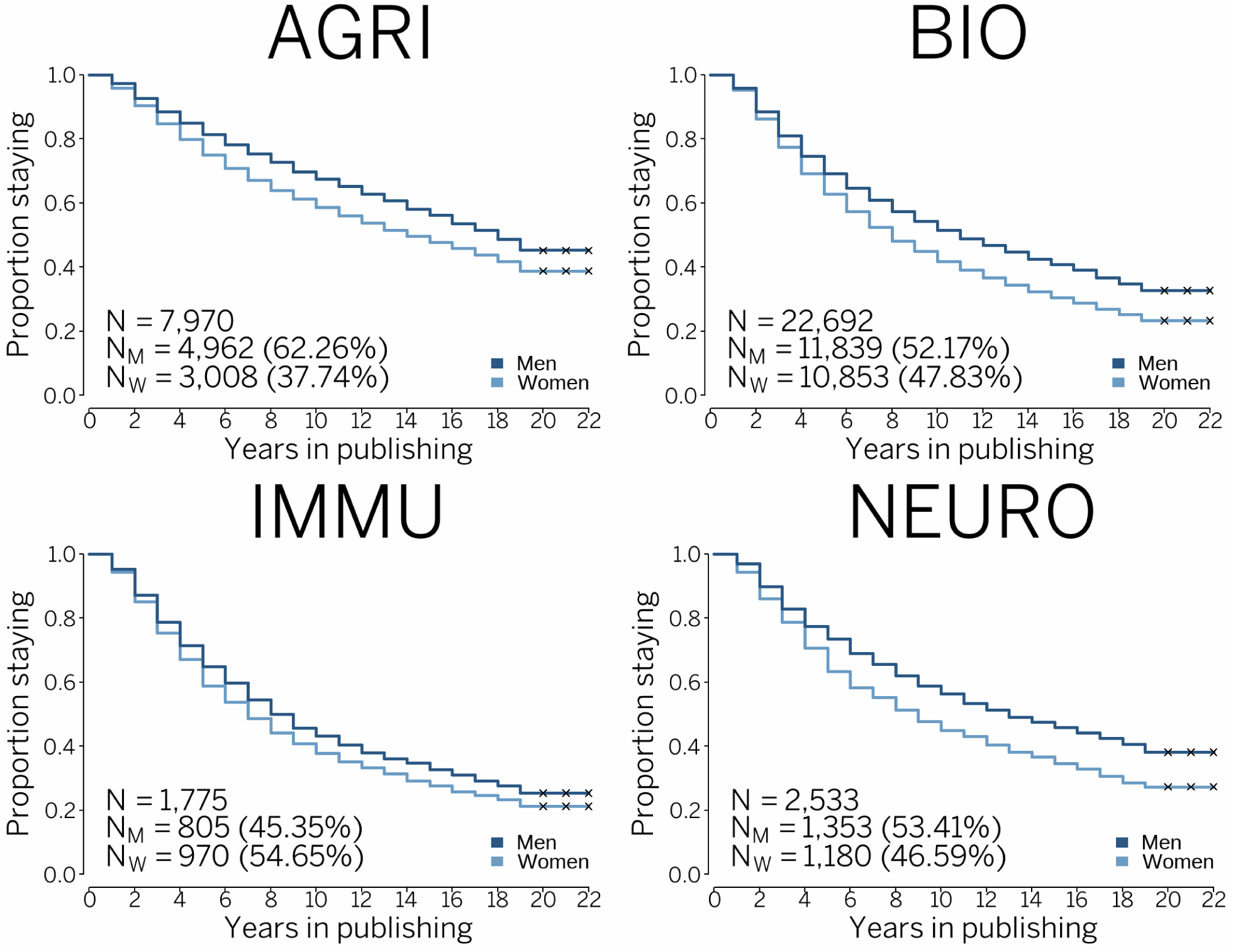
2000 cohort of scientists, Kaplan‒Meier survival curve, AGRI, agricultural and biological sciences (N=7,970), BIO, biochemistry, genetics, and molecular biology (N=22,692), IMMU, immunology and microbiology (N=1,775), and NEURO, neuroscience (N=2,533). The four disciplines combined (N=34,970).

For the 2000 cohort, we can observe, year after year, how the probability of staying in science is always higher for men and always lower for women. Similarly, Figure 2 shows gender differences for the 2010 cohort. What is clearly visible at the level of survival curves is that attrition levels for the younger cohort of scientists are higher (and consequently retention levels are lower), both for men and women, than for the older cohort. Young people, usually in the career stage of doctoral school, come to biology and disappear from it much faster in the younger cohort. At the same time, the two pictures of the two cohorts coming and staying in science differ considerably in gender terms. The differences for the younger cohort are lower, especially in the two largest disciplines examined, namely AGRI and BIO.

**Figure 2.**
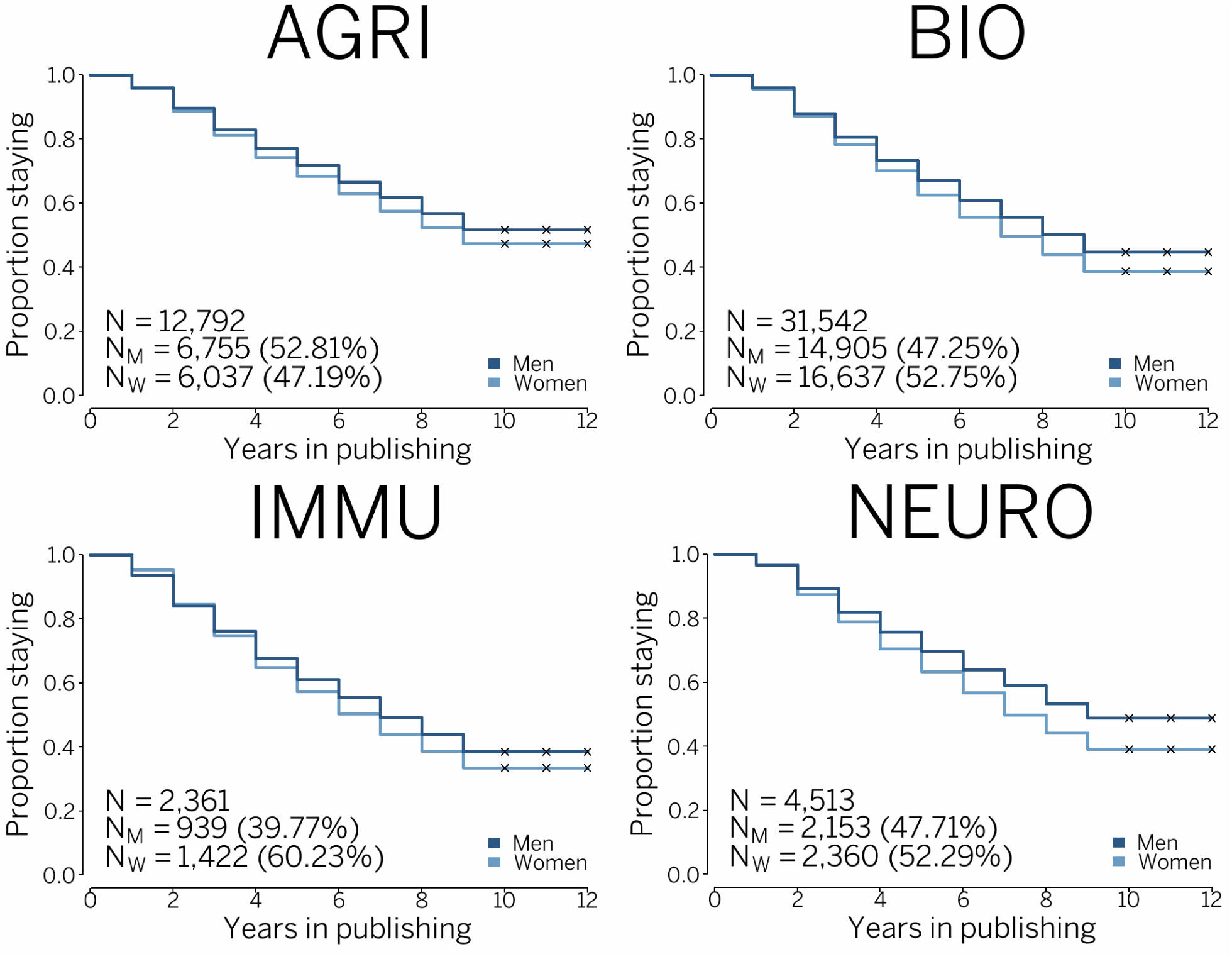
2010 cohort of scientists, Kaplan‒Meier survival curve, AGRI, agricultural and biological sciences (N=12,792), BIO, biochemistry, genetics, and molecular biology (N=31,542), IMMU, immunology and microbiology (N=2,361), and NEURO, neuroscience (N=4,513). The four disciplines combined (N=51,208).

None of the disciplines was resistant to changes from a temporal perspective. Time matters in science: while there were ever more women in the disciplines (Table 1), gender differences in the attrition (and retention) rates became smaller over time.

Overall, 62.8% of women in the 2000 cohort in BIO were still in science after 5 years, 41.7% after 10 years, and only 23.4% by the end of the period examined (or after 19 years). For men in BIO, the percentages were substantially higher: 69.2%, 51.4%, and 32.7%, respectively (Table 2). These numbers highlight how differently, on average, male and female careers developed over time.

**Table 2:** 2000 and 2010 cohorts of scientists, BIO, biochemistry, genetics, and molecular biology. Kaplan–Meier estimate by gender with total counts for men and women, time (in years), number of observations of scientists leaving science. Kaplan–Meier probability of staying in science with a 95% confidence interval. Note: (1) standard error of 0.01. (Total i.e., men and women combined in Supplementary Table 1).

| Time (years) | Women (cohort 2000) |  |  | Men (cohort 2000) |  |  | Probability of staying is higher for men by ... (in %) |
| --- | --- | --- | --- | --- | --- | --- | --- |
|  | n | n leaving science | KM probability (staying) with 95% CI and SE | n | n leaving science | KM probability (staying) with 95% CI and SE |  |
| 1 | 10,853 | 526 | 0.952 (0.948-0.956)<sup>1</sup> | 11,839 | 494 | 0.958 (0.958-0.958)<sup>1</sup> | **0.63** |
| 2 | 10,327 | 971 | 0.862 (0.856-0.869)<sup>1</sup> | 11,345 | 865 | 0.885 (0.885-0.885)<sup>1</sup> | **2.67** |
| 3 | 9,356 | 956 | 0.774 (0.766-0.782)<sup>1</sup> | 10,480 | 900 | 0.809 (0.809-0.809)<sup>1</sup> | **4.52** |
| 4 | 8,400 | 901 | 0.691 (0.682-0.700)<sup>1</sup> | 9,580 | 754 | 0.746 (0.746-0.746)<sup>1</sup> | **7.96** |
| 5 | 7,499 | 678 | 0.628 (0.619-0.638)<sup>1</sup> | 8,826 | 637 | 0.692 (0.692-0.692)<sup>1</sup> | **10.19** |
| 6 | 6,821 | 607 | 0.573 (0.563-0.582)<sup>1</sup> | 8,189 | 530 | 0.647 (0.647-0.647)<sup>1</sup> | **12.91** |
| 7 | 6,214 | 526 | 0.524 (0.515-0.534)<sup>1</sup> | 7,659 | 445 | 0.609 (0.609-0.609)<sup>1</sup> | **16.22** |
| 8 | 5,688 | 465 | 0.481 (0.472-0.491)<sup>1</sup> | 7,214 | 435 | 0.573 (0.573-0.573)<sup>1</sup> | **19.13** |
| 9 | 5,223 | 353 | 0.449 (0.439-0.458)<sup>1</sup> | 6,779 | 349 | 0.543 (0.543-0.543)<sup>1</sup> | **20.94** |
| 10 | 4,870 | 345 | 0.417 (0.408-0.426)<sup>1</sup> | 6,430 | 341 | 0.514 (0.514-0.514)<sup>1</sup> | **23.26** |
| 11 | 4,525 | 290 | 0.390 (0.381-0.400)<sup>1</sup> | 6,089 | 292 | 0.490 (0.490-0.490)<sup>1</sup> | **25.64** |
| 12 | 4,235 | 255 | 0.367 (0.358-0.376)<sup>1</sup> | 5,797 | 254 | 0.468 (0.468-0.468)<sup>1</sup> | **27.52** |
| 13 | 3,980 | 249 | 0.344 (0.335-0.353)<sup>1</sup> | 5,543 | 255 | 0.447 (0.447-0.447)<sup>1</sup> | **29.94** |
| 14 | 3,731 | 210 | 0.324 (0.316-0.333)<sup>1</sup> | 5,288 | 251 | 0.425 (0.425-0.425)<sup>1</sup> | **31.17** |
| 15 | 3,521 | 212 | 0.305 (0.296-0.314)<sup>1</sup> | 5,037 | 212 | 0.408 (0.408-0.408)<sup>1</sup> | **33.77** |
| 16 | 3,309 | 189 | 0.287 (0.279-0.296)<sup>1</sup> | 4,825 | 205 | 0.390 (0.390-0.390)<sup>1</sup> | **35.89** |
| 17 | 3,120 | 198 | 0.269 (0.261-0.278)<sup>1</sup> | 4,620 | 279 | 0.367 (0.367-0.367)<sup>1</sup> | **36.43** |
| 18 | 2,922 | 184 | 0.252 (0.244-0.261)<sup>1</sup> | 4,341 | 221 | 0.348 (0.348-0.348)<sup>1</sup> | **38.10** |
| 19 | 2,738 | 203 | 0.234 (0.226-0.242)<sup>1</sup> | 4,120 | 243 | 0.327 (0.327-0.327)<sup>1</sup> | **39.74** |
| Time (years) | Women (cohort 2010) |  |  | Men (cohort 2010) |  |  | Probability of staying is higher for men by ... (in %) |
|  | n | n leaving science | KM probability (staying) with 95% CI and SE | n | n leaving science | KM probability (staying) with 95% CI and SE |  |
| 1 | 16,637 | 715 | 0.957 (0.954-0.960)<sup>1</sup> | 14,905 | 600 | 0.960 (0.960-0.960)<sup>1</sup> | **0.31** |
| 2 | 15,922 | 1,423 | 0.871 (0.866-0.877)<sup>1</sup> | 14,305 | 1,199 | 0.879 (0.879-0.879)<sup>1</sup> | **0.92** |
| 3 | 14,499 | 1,464 | 0.783 (0.777-0.790)<sup>1</sup> | 13,106 | 1,085 | 0.807 (0.807-0.807)<sup>1</sup> | **3.07** |
| 4 | 13,035 | 1,389 | 0.700 (0.693-0.707)<sup>1</sup> | 12,021 | 1,092 | 0.733 (0.733-0.733)<sup>1</sup> | **4.71** |
| 5 | 11,646 | 1,246 | 0.625 (0.618-0.633)<sup>1</sup> | 10,929 | 940 | 0.670 (0.670-0.670)<sup>1</sup> | **7.20** |
| 6 | 10,400 | 1,158 | 0.556 (0.548-0.563)<sup>1</sup> | 9,989 | 901 | 0.610 (0.610-0.610)<sup>1</sup> | **9.71** |
| 7 | 9,242 | 980 | 0.497 (0.489-0.504)<sup>1</sup> | 9,088 | 805 | 0.556 (0.556-0.556)<sup>1</sup> | **11.87** |
| 8 | 8,262 | 931 | 0.441 (0.433-0.448)<sup>1</sup> | 8,283 | 806 | 0.502 (0.502-0.502)<sup>1</sup> | **13.83** |
| 9 | 7,331 | 895 | 0.387 (0.380-0.394)<sup>1</sup> | 7,477 | 797 | 0.448 (0.448-0.448)<sup>1</sup> | **15.76** |

Thus, women in BIO were 10.19% more likely to drop out of science than men after 5 years, with markedly increasing chances of dropping out in later years. Women were 23.26% more likely to leave science after 10 years, and 39.74% more likely to leave science before the end of the studied period. Gender difference in attrition, only slightly visible after 5 years, consistently increased in later career stages.

Similarly, women in NEURO were 15.77% more likely to drop out than men after 5 years, 25.61% more likely to leave science after 10 years, and 39.42% more likely to leave science by the end of the studied period. After 19 years (i.e., only uncensored observations), the chances of surviving in either BIO or NEURO were 40% higher for men than for women (Table 2 and Supplementary Table 4). In the two other disciplines, gender differences in attrition were considerably lower. For AGRI, the chances of surviving 10 years were 15.21% higher for men, and 19 years, 16.49% higher for men (Table 3). For IMMU, our data show 14.29% and 20.28%, respectively (Supplementary Table 3).

**Table 3:** 2000 and 2010 cohorts of scientists, AGRI, agricultural and biological sciences. Kaplan–Meier estimate by gender with total counts for men and women, time (in years), number of observations of scientists leaving science. Kaplan–Meier probability of staying in science with a 95% confidence interval. Note: (1) standard error of 0.01. (Total i.e., men and women combined in Supplementary Table 2).

| Time (years) | Women (cohort 2000) |  |  | Men (cohort 2000) |  |  | Probability of staying is higher for men by ... (in %) |
| --- | --- | --- | --- | --- | --- | --- | --- |
|  | n | n leaving science | KM probability (staying) with 95% CI and SE | n | n leaving science | KM probability (staying) with 95% CI and SE |  |
| 1 | 3,008 | 125 | 0.958 (0.951-0.966)<sup>1</sup> | 4,962 | 138 | 0.972 (0.972-0.972)<sup>1</sup> | **1.46** |
| 2 | 2,883 | 167 | 0.903 (0.892-0.914)<sup>1</sup> | 4,824 | 226 | 0.927 (0.927-0.927)<sup>1</sup> | **2.66** |
| 3 | 2,716 | 170 | 0.846 (0.834-0.859)<sup>1</sup> | 4,598 | 212 | 0.884 (0.884-0.884)<sup>1</sup> | **4.49** |
| 4 | 2,546 | 145 | 0.798 (0.784-0.813)<sup>1</sup> | 4,386 | 169 | 0.850 (0.850-0.850)<sup>1</sup> | **6.52** |
| 5 | 2,401 | 144 | 0.750 (0.735-0.766)<sup>1</sup> | 4,217 | 182 | 0.813 (0.813-0.813)<sup>1</sup> | **8.40** |
| 6 | 2,257 | 125 | 0.709 (0.693-0.725)<sup>1</sup> | 4,035 | 158 | 0.781 (0.781-0.781)<sup>1</sup> | **10.16** |
| 7 | 2,132 | 115 | 0.671 (0.654-0.688)<sup>1</sup> | 3,877 | 142 | 0.753 (0.753-0.753)<sup>1</sup> | **12.22** |
| 8 | 2,017 | 95 | 0.639 (0.622-0.656)<sup>1</sup> | 3,735 | 131 | 0.726 (0.726-0.726)<sup>1</sup> | **13.62** |
| 9 | 1,922 | 82 | 0.612 (0.595-0.629)<sup>1</sup> | 3,604 | 146 | 0.697 (0.697-0.697)<sup>1</sup> | **13.89** |
| 10 | 1,840 | 79 | 0.585 (0.568-0.603)<sup>1</sup> | 3,458 | 113 | 0.674 (0.674-0.674)<sup>1</sup> | **15.21** |
| 11 | 1,761 | 76 | 0.560 (0.543-0.578)<sup>1</sup> | 3,345 | 113 | 0.651 (0.651-0.651)<sup>1</sup> | **16.25** |
| 12 | 1,685 | 70 | 0.537 (0.519-0.555)<sup>1</sup> | 3,232 | 115 | 0.628 (0.628-0.628)<sup>1</sup> | **16.95** |
| 13 | 1,615 | 65 | 0.515 (0.498-0.533)<sup>1</sup> | 3,117 | 109 | 0.606 (0.606-0.606)<sup>1</sup> | **17.67** |
| 14 | 1,550 | 59 | 0.496 (0.478-0.514)<sup>1</sup> | 3,008 | 124 | 0.581 (0.581-0.581)<sup>1</sup> | **17.14** |
| 15 | 1,491 | 53 | 0.478 (0.461-0.496)<sup>1</sup> | 2,884 | 97 | 0.562 (0.562-0.562)<sup>1</sup> | **17.57** |
| 16 | 1,438 | 59 | 0.458 (0.441-0.477)<sup>1</sup> | 2,787 | 126 | 0.536 (0.536-0.536)<sup>1</sup> | **17.03** |
| 17 | 1,379 | 60 | 0.438 (0.421-0.457)<sup>1</sup> | 2,661 | 104 | 0.515 (0.515-0.515)<sup>1</sup> | **17.58** |
| 18 | 1,319 | 61 | 0.418 (0.401-0.436)<sup>1</sup> | 2,557 | 144 | 0.486 (0.486-0.486)<sup>1</sup> | **16.27** |
| 19 | 1,258 | 90 | 0.388 (0.371-0.406)<sup>1</sup> | 2,413 | 169 | 0.452 (0.452-0.452)<sup>1</sup> | **16.49** |
| Time (years) | Women (cohort 2010) |  |  | Men (cohort 2010) |  |  | Probability of staying is higher for men by ... (in %) |
|  | n | n leaving science | KM probability (staying) with 95% CI and SE | n | n leaving science | KM probability (staying) with 95% CI and SE |  |
| 1 | 6,037 | 252 | 0.958 (0.953-0.963)<sup>1</sup> | 6,755 | 265 | 0.961 (0.961-0.961)<sup>1</sup> | **0.31** |
| 2 | 5,785 | 431 | 0.887 (0.879-0.895)<sup>1</sup> | 6,490 | 442 | 0.895 (0.895-0.895)<sup>1</sup> | **0.90** |
| 3 | 5,354 | 456 | 0.811 (0.802-0.821)<sup>1</sup> | 6,048 | 452 | 0.828 (0.828-0.828)<sup>1</sup> | **2.10** |
| 4 | 4,898 | 415 | 0.743 (0.732-0.754)<sup>1</sup> | 5,596 | 397 | 0.770 (0.770-0.770)<sup>1</sup> | **3.63** |
| 5 | 4,483 | 351 | 0.684 (0.673-0.696)<sup>1</sup> | 5,199 | 351 | 0.718 (0.718-0.718)<sup>1</sup> | **4.97** |
| 6 | 4,132 | 330 | 0.630 (0.618-0.642)<sup>1</sup> | 4,848 | 354 | 0.665 (0.665-0.665)<sup>1</sup> | **5.56** |
| 7 | 3,802 | 331 | 0.575 (0.563-0.588)<sup>1</sup> | 4,494 | 317 | 0.618 (0.618-0.618)<sup>1</sup> | **7.48** |
| 8 | 3,471 | 305 | 0.524 (0.512-0.537)<sup>1</sup> | 4,177 | 342 | 0.568 (0.568-0.568)<sup>1</sup> | **8.40** |
| 9 | 3,166 | 311 | 0.473 (0.460-0.486)<sup>1</sup> | 3,835 | 338 | 0.518 (0.518-0.518)<sup>1</sup> | **9.51** |

The Kaplan–Meier survival curves reflect the underlying data produced for each cohort, year, discipline, and gender. Table 2 (for BIO) and Table 3 (for AGRI) highlight the changing Kaplan–Meier probability of staying in science (with a 95% confidence interval) separately for women and men in each year starting with 2000 (or 2010).

Taking scientists in BIO as an example, in year 2000, 10,853 women and 11,839 men had their first publication, but in year 1 (Table 2), 526 women and 494 men stopped publishing. In year 1, the difference in retention (and, by definition, in attrition, its mirror image) between men and women was small but statistically significant. However, the gender difference in retention increased with every year, from 10.19% after 5 years to 39.74% after 19 years. Gender differences were substantial and statistically significant, with narrow confidence intervals indicating high concentrations of observations around the percentages shown.

In the case of the 2010 BIO cohort, the gender difference in retention was lower, and after 9 years, that is, in the last year examined in our study, 38.7% of women stayed in science, as opposed to 44.8% of men. For this younger cohort, the probability of staying in science after 9 years was 15.76% higher for men, compared with 20.94% in the older cohort.

Importantly, while gender differences in attrition diminished, attrition became an ever more acute problem: after 9 years (with no data beyond this period), retention rates decreased by more than 15 percentage points (from 61.2% to 47.3% for women and from 69.7% to 51.8% for men). The chances of surviving in science became smaller for both men and women. This indicates an issue requiring reflection not only on women but on the science workforce in general.

Comparing the four disciplines, for the older cohort, gender differences in retention were highest for NEURO after 5 years, after 10 years, and at end of the period examined, reaching 10% in each case. Cross-disciplinary differences were marked, with only about 20% of women in IMMU contrasted with about 40% in AGRI remaining in science at the end of the period.

Comparing the four disciplines for the younger cohort after 9 years, the highest probability of staying in science was observed for AGRI (47.3% for women, 51.8% for men), with women in the other disciplines having probabilities below 40%. The gender difference was highest for NEURO, reaching nearly 10 percentage points (39.1% vs. 48.9%).

The estimated probability that a woman would survive in science 15 years differed considerably between the four disciplines: it varied from as much as 47.8% in AGRI (95% confidence interval: 0.461–0.496) to as little as 27.6%, or 20 p.p. less, in IMMU (0.250– 0.306). For men, the probabilities were much higher at 56.2% and 32.7% (higher by 17.57% and 18.48%, respectively; Table 2 and Supplementary Table 2).

Our data show that women in BIO (as well as those AGRI and NEURO) disappeared from science, with the passage of time, in ever-larger proportions compared with men. In contrast, women in male-dominated disciplines – with shares of women in the range of 15%–20%, namely PHYS, COMP, and MATH, disappeared from science in almost exactly the same proportions as men – with the two Kaplan-Meier curves overlapping and no statistically significant differences – in the entire period examined (Kwiek & Szymula, 2024).

For a more comprehensive view of gender differences in attrition and retention in the four disciplines, we correlated the data on both cohorts to discuss three other approaches – survival regression curves, hazard rate curves, and kernel density curves – separately for each discipline.

Survival analysis models the time until an event of interest (here, ceasing publishing) occurs. For some proportion of the 2000 and 2010 cohorts, the event occurred. Hazard is the probability of an event occurring during any given time point within a period. We calculated the probability that any given scientist would cease publishing each year and plotted the results. The survival regression curve is actually a smoothed Kaplan‒Meier curve. It starts with 1 on a y-axis because all scientists are publishing at time zero, then gradually, the share of authors declines.

The survival regression curves for the four disciplines for both cohorts (Figure 3, Figure 4 Panels B; as well as Supplementary Figures 1-2) indicate a steeper decline for men and women in the early years of their publishing careers and a smoother decline in later years, with increasing divergence between the curves for men and women from the 2000 cohort.

**Figure 3:**
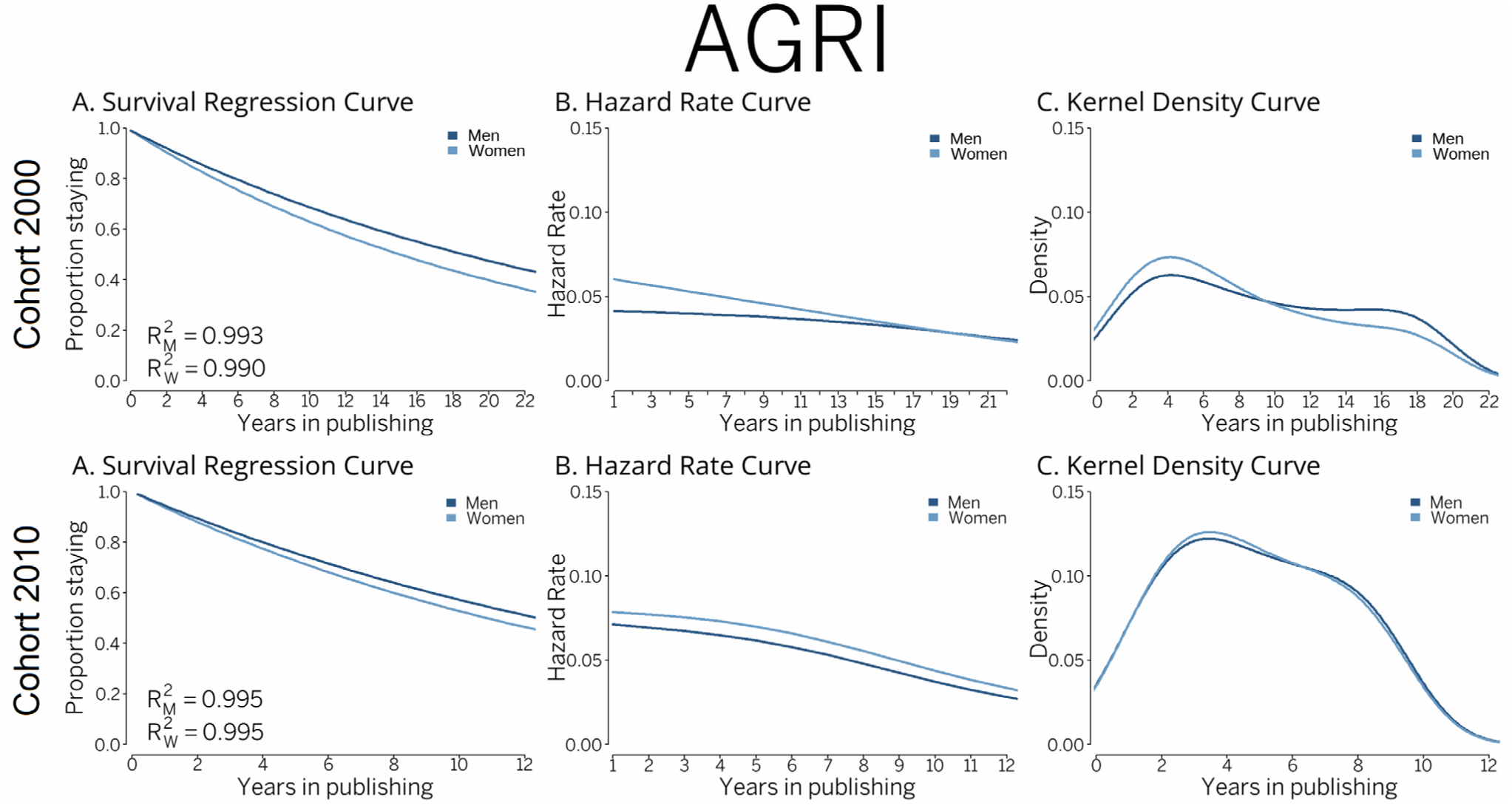
2000 (top panel, N=7,970) and 2010 (bottom panel, N=12,792) cohorts of scientists, AGRI, agricultural and biological sciences. Survival regression curve (exponential distribution fitting of Kaplan‒Meier curve), hazard rate curve (B-splines smoothing method, smoothing parameter 10k), and kernel density curve (B-splines smoothing method, bandwidth 2, component per point based on Gaussian curve)

**Figure 4:**
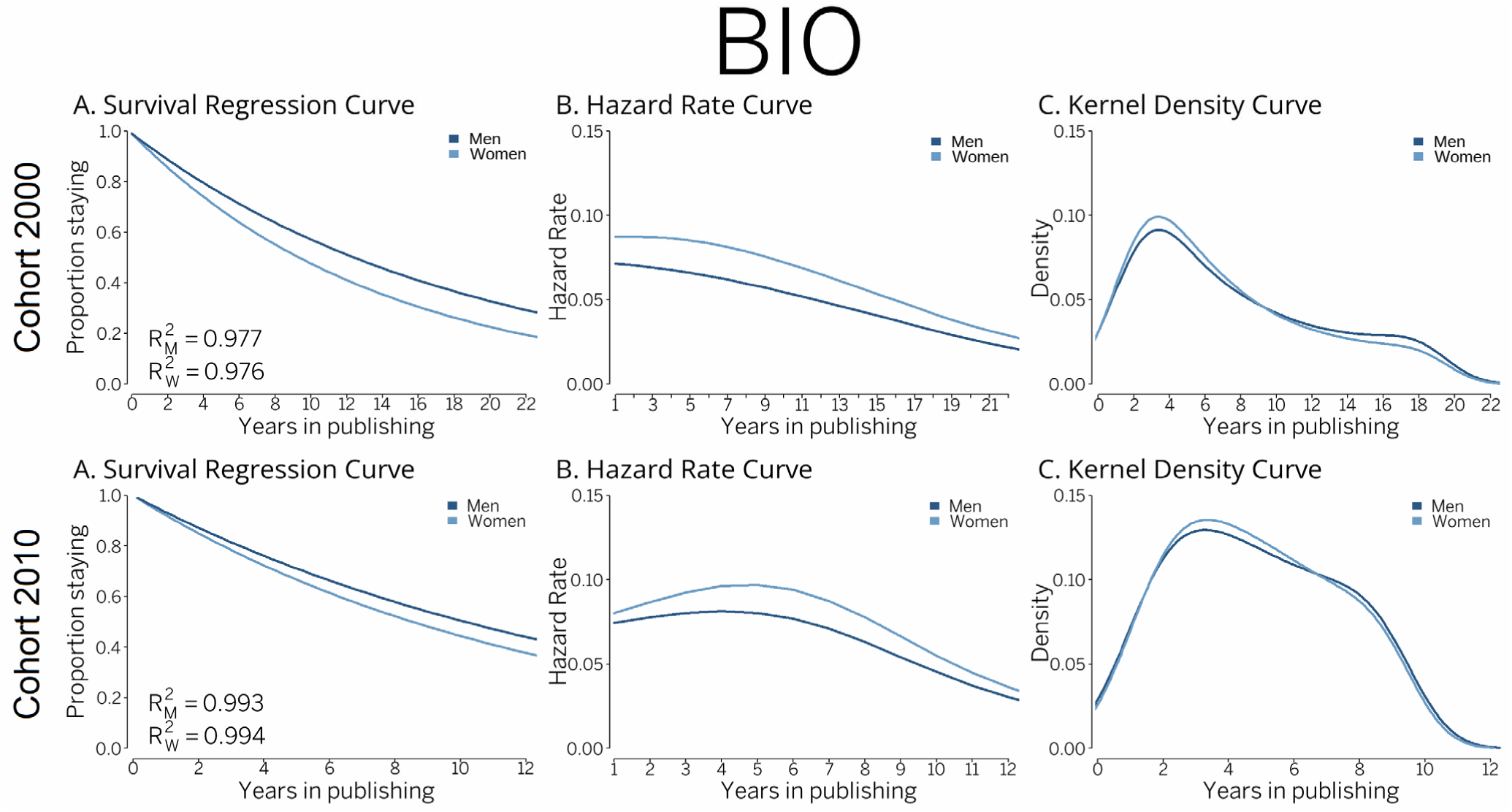
2000 (top panel, N=22,692) and 2010 (bottom panel, N=31,542) cohorts of scientists, BIO, biochemistry, genetics, and molecular biology. Survival regression curve (exponential distribution fitting of Kaplan‒Meier curve), hazard rate curve (B-splines smoothing method, smoothing parameter 10k), and kernel density curve (B-splines smoothing method, bandwidth 2, component per point based on Gaussian curve)

The hazard rate curves (panels C) show that attrition was greater in the early publishing years. Especially in BIO and NEURO for the 2000 cohort and for the first 15 years, women had greater chances of leaving science than men. The hazard functions shown for the 2000 cohort monotonically decrease: The short-term risk of ceasing publishing is greater in the early years of publishing careers, after which the risk decreases over time. The riskiest years are always the early years in science. Most importantly, gender differences are visible for all disciplines examined, with the curves for men and women never overlapping (except for AGRI, cohort 2000).

Kernel density curves (Figure 4, Panels D) use kernel density estimation to create a smoothed, continuous curve that approximates the underlying data distribution. Kernel density estimations display the distribution of values in a dataset using one continuous curve identifying the shape of the distribution. Kernel density curves show how all men and women who actually left science are distributed over time. The curves are easily interpretable because the area under the curve always reaches 100%. For both cohorts, the shares of scientists who left science were highest for the first 2–4 years, with peaks in year 4. This likely represents the doctoral stage, which does not always lead to postdoctoral or full employment.

The survival regression curves mirror the Kaplan–Meier curves. They highlight consistent and large gender differences in attrition, especially in BIO and NEURO. The differences visible for the older cohort are still visible for the younger cohort. The hazard rate curves show that in two disciplines (NEURO and IMMU), the chances of leaving science were as gendered for the younger cohort as they were for the older one. Finally, kernel density curves show largely the same stories about the four disciplines: for both cohorts, disappearing from science was more intensive in the first 4 years, and much less intensive after 10 years. The distribution patterns of the leavers were largely similar for men and women.

## Conclusions

The conceptualization of stopping publishing as leaving science does not entail any other academic or non-academic roles (teaching and administration, as well as working for governments, NGOs, industry, etc.). In our simplified methodological approach, a sequence of globally indexed publications replaces a sequence of much wider cognitive and social processes encompassing the various dimensions of ‘doing science’ (Sugimoto & Larivière, 2023; Kwiek, 2019). In it, not publishing in scholarly journals means not doing science anymore – which is a fairly simple approach driven by data availability for a global study like ours.

The metadata about publications at an individual level, with considerable cleaning and computing, are globally (and longitudinally) available for comparative research. In contrast, micro-data about employment in the science sector – first employment, changing sectors – including academia, are not available. Our research would not be possible without access to micro-data drawn from the raw Scopus dataset and computed using commercial, cloud-based computing infrastructure.

Our study has an OECD focus: all data refer to 38 OECD countries combined. However, it is also possible to compare differences by country. We have published an interactive tool that provides the results of the Kaplan–Meier probability of staying in science by country, discipline, and gender for 11 cohorts (the 2000–2010 cohorts, N=2,127,803). The snapshot in Figure 5 shows the probability of staying in science after 10 years for women from the 2000 cohort for major European countries in biochemistry, genetics, and molecular biology (BIO). It reaches only 30% in Germany, as opposed to the much higher 83% in Portugal and 64% in Poland.

**Figure 5.**
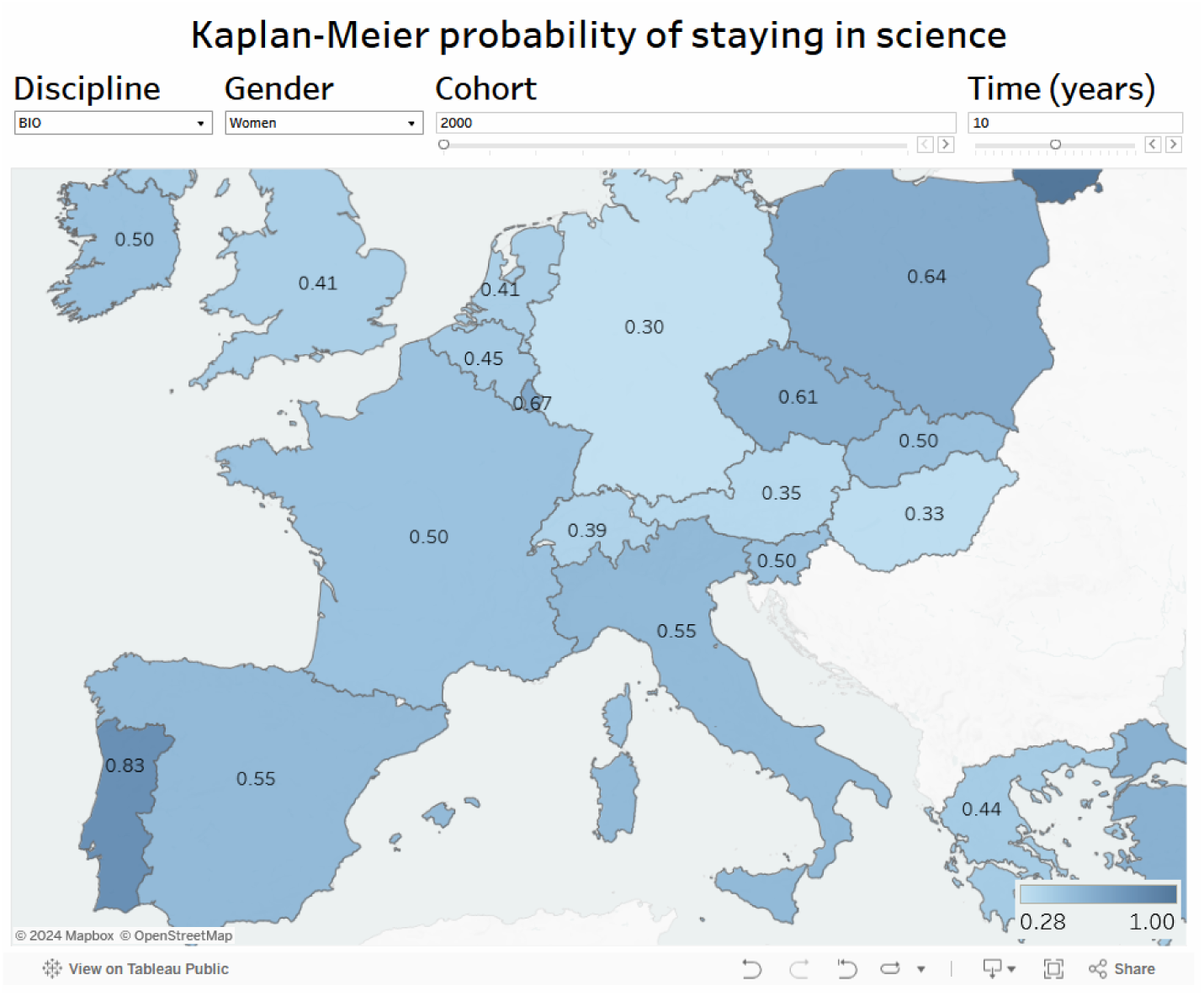
BIO, biochemistry, genetics, and molecular biology. Women scientists from cohort 2000, probability of staying in science after 10 years. A snapshot of an interactive map (available at https://public.tableau.com/app/profile/marek.kwiek/viz/Attrition-in-science-OECD/Dashboard) in which Kaplan‒Meier probabilities of remaining in science (i.e., continuing publishing) are provided by country (38 OECD countries), discipline, and gender and for all 11 cohorts from 2000–2010 (N=2,127,803 scientists in 16 STEMM disciplines, of which 1,289,756 are identified as men and 838,047 as women).

The chances of surviving in biochemistry, genetics, and molecular biology greatly differ across Europe. The various retention rates reflect different national conditions in the science sector changing over time. In some countries, push factors are important in leaving decisions (e.g., the science sector being an unattractive, perhaps unwelcoming workplace for women), while in others, pull factors matter (e.g., outside opportunities are attractive, pulling scientists to industry). Examples include computing in the USA, chemistry and engineering in Germany, with attractive non-academic workplaces – as opposed to almost all disciplines in Poland and Portugal. There, pull factors have traditionally been weak, with outside, non-academic opportunities for scientists being limited.

Exploring attrition and retention in science at a multi-country scale has limitations – but it also creates new opportunities. Disciplines can be examined across many countries, and conclusions can be drawn about cross-disciplinary differences within the STEMM disciplines. The traditional “leaky pipeline” and “chilly climate” metaphors can be tested in detail, using large populations of scientists from different countries and different disciplines. While regarded as a monolith, there are large variations among STEMM disciplines, especially in their gender composition over time. With global datasets, attrition and retention – coming to science and leaving it – can be explored from a longitudinal perspective in which the same individual scientists are tracked for years and decades, rather than from the perspective of annually changing averages in the science profession by gender in countries and institutions. These are known from traditional science and engineering statistics at an aggregated level, which hide the different career trajectories of men and women.

Attrition, as it emerged from our results for scientists in AGRI, BIO, IMMU, and NEURO, is a permanent challenge in science. Attrition affects men and women alike – but in these disciplines, with gender balance already achieved (according to the 40/60 formula), attrition affects women much more powerfully than men.

About 60% of women from the 2000 cohort in BIO were still in science after 5 years, 40% after 10 years, and only 20% at the end of the period examined (i.e., after 19 years). In stark contrast, the percentages were substantially higher for men: approximately 70%, 50%, and 30%, respectively (Table 2). These reflect enormous differences in how, on average, male and female careers develop over time. Kaplan–Meier estimations show that women in BIO were one quarter more likely to leave science after 10 years (23.26%), and as much as 40% more likely to leave science by the end of the study period (39.74%). The data throw new light on careers in BIO (and related fields) in the 38 OECD countries combined.

Our online interactive tool (Figure 5) reports data about retention in BIO for the USA (the largest OECD science system): 60% vs. 70% after 5 years, 40% vs. 51% after 10 years, and 20% vs. 30% at the end of the study period for women vs. men, respectively. The Kaplan– Meier probability of surviving 19 years in BIO was 50% higher for men (61.11% higher for men in IMMU, 52% in NEURO, but only 20.59% in AGRI – for which the highest retention rates were also observable for each year).

It is intriguing that in the disciplines where gender balance has been achieved (as in the four studied here), for both older and younger cohorts of scientists, the chances of women surviving were consistently lower than for men. However, in disciplines where there is a long way to go to achieve equal representation of men and women (as in mathematics, computing, or engineering), for both cohorts, women’s chances of surviving were equal to those of men, with no statistically significant differences at all.

Consequently, increasing the representation of women in science does not automatically guarantee increasingly equal attrition rates for men and women. In short, the problem for women in male-dominated disciplines is how to get there (conceptualized with our dataset as how to start publishing). The problem for women in the biology-related disciplines examined here, with equal gender representation, is not so much getting there – it is actually staying there (i.e., continuing to publish) for as long as men do.

Our research shows that attrition means different things for men and women in different STEMM disciplines, and different things for men and women belonging to different cohorts. In the four disciplines examined here, gender differences in attrition and retention have been and continue to be high and are only slightly decreasing over time.

*This paper is accompanied by* Electronic Supplementary Material, *which is available online*.

## Acknowledgments

MK gratefully acknowledges the support provided by MNISW NDS grant no. NdS-II/SP/0010/2023/01. LS is grateful for the support of his doctoral studies provided by NCN grant 2019/35/0/HS6/02591. We gratefully acknowledge the assistance of the International Center for the Studies of Research (ICSR) Lab with Kristy James and Alick Bird. We also want to thank Dr. Wojciech Roszka from the CPPS Poznan Team for many fruitful discussions.

---

**Supplementary Table 1:** 2000 and 2010 cohorts of scientists, BIO, biochemistry, genetics, and molecular biology. Kaplan–Meier estimate with total counts (men and women combined), time (in years), number of observations of scientists leaving science. Kaplan–Meier probability of staying in science with a 95% confidence interval. Note: (1) standard error of 0.01.

| Total (cohort 2000) |  |  |  |
| --- | --- | --- | --- |
| Time (years) | n | n leaving science | KM probability (staying) with 95% CI and SE |
| 1 | 22,692 | 1,020 | 0.955 (0.951–0.958)<sup>1</sup> |
| 2 | 21,672 | 1,836 | 0.866 (0.861–0.870)<sup>1</sup> |
| 3 | 19,836 | 1,856 | 0.778 (0.772–0.783)<sup>1</sup> |
| 4 | 17,980 | 1,655 | 0.704 (0.698–0.710)<sup>1</sup> |
| 5 | 16,325 | 1,315 | 0.645 (0.639–0.652)<sup>1</sup> |
| 6 | 15,010 | 1,137 | 0.592 (0.586–0.598)<sup>1</sup> |
| 7 | 13,873 | 971 | 0.543 (0.537–0.550)<sup>1</sup> |
| 8 | 12,902 | 900 | 0.500 (0.494–0.507)<sup>1</sup> |
| 9 | 12,002 | 702 | 0.466 (0.459–0.472)<sup>1</sup> |
| 10 | 11,300 | 686 | 0.434 (0.427–0.441)<sup>1</sup> |
| 11 | 10,614 | 582 | 0.408 (0.401–0.414)<sup>1</sup> |
| 12 | 10,032 | 509 | 0.385 (0.378–0.392)<sup>1</sup> |
| 13 | 9,523 | 504 | 0.363 (0.356–0.370)<sup>1</sup> |
| 14 | 9,019 | 461 | 0.344 (0.337–0.351)<sup>1</sup> |
| 15 | 8,558 | 424 | 0.325 (0.318–0.332)<sup>1</sup> |
| 16 | 8,134 | 394 | 0.308 (0.301–0.315)<sup>1</sup> |
| 17 | 7,740 | 477 | 0.288 (0.281–0.295)<sup>1</sup> |
| 18 | 7,263 | 405 | 0.271 (0.264–0.278)<sup>1</sup> |
| 19 | 6,858 | 446 | 0.253 (0.246–0.260)<sup>1</sup> |
| Total (cohort 2010) |  |  |  |
| 1 | 31,542 | 1,315 | 0.959 (0.956–0.961)<sup>1</sup> |
| 2 | 30,227 | 2,622 | 0.875 (0.871–0.879)<sup>1</sup> |
| 3 | 27,605 | 2,549 | 0.795 (0.790–0.800)<sup>1</sup> |
| 4 | 25,056 | 2,481 | 0.717 (0.712–0.722)<sup>1</sup> |
| 5 | 22,575 | 2,186 | 0.652 (0.647–0.658)<sup>1</sup> |
| 6 | 20,389 | 2,059 | 0.590 (0.584–0.596)<sup>1</sup> |
| 7 | 18,330 | 1,785 | 0.534 (0.528–0.540)<sup>1</sup> |
| 8 | 16,545 | 1,737 | 0.478 (0.472–0.484)<sup>1</sup> |
| 9 | 14,808 | 1,692 | 0.418 (0.412–0.424)<sup>1</sup> |

**Supplementary Table 2:** 2000 and 2010 cohorts of scientists, AGRI, agricultural and biological sciences. Kaplan– Meier estimate with total counts (men and women combined), time (in years), number of observations of scientists leaving science. Kaplan–Meier probability of staying in science with a 95% confidence interval. Note: (1) standard error of 0.01.

| **Total (cohort 2000)** |  |  |  |
| --- | --- | --- | --- |
| **Time (years)** | **n** | **n leaving science** | **KM probability (staying) with 95% CI and SE** |
| 1 | 7,970 | 263 | 0.967 (0.964–0.970)<sup>1</sup> |
| 2 | 7,707 | 393 | 0.917 (0.911–0.923)<sup>1</sup> |
| 3 | 7,314 | 382 | 0.865 (0.859–0.872)<sup>1</sup> |
| 4 | 6,932 | 314 | 0.826 (0.819–0.834)<sup>1</sup> |
| 5 | 6,618 | 326 | 0.783 (0.776–0.791)<sup>1</sup> |
| 6 | 6,292 | 283 | 0.749 (0.741–0.757)<sup>1</sup> |
| 7 | 6,009 | 257 | 0.718 (0.710–0.726)<sup>1</sup> |
| 8 | 5,752 | 226 | 0.689 (0.681–0.698)<sup>1</sup> |
| 9 | 5,526 | 228 | 0.659 (0.650–0.668)<sup>1</sup> |
| 10 | 5,298 | 192 | 0.632 (0.623–0.641)<sup>1</sup> |
| 11 | 5,106 | 189 | 0.607 (0.598–0.616)<sup>1</sup> |
| 12 | 4,917 | 185 | 0.584 (0.575–0.593)<sup>1</sup> |
| 13 | 4,732 | 174 | 0.561 (0.552–0.570)<sup>1</sup> |
| 14 | 4,558 | 183 | 0.540 (0.531–0.550)<sup>1</sup> |
| 15 | 4,375 | 150 | 0.521 (0.512–0.531)<sup>1</sup> |
| 16 | 4,225 | 185 | 0.498 (0.489–0.507)<sup>1</sup> |
| 17 | 4,040 | 164 | 0.477 (0.468–0.487)<sup>1</sup> |
| 18 | 3,876 | 205 | 0.449 (0.440–0.459)<sup>1</sup> |
| 19 | 3,671 | 259 | 0.413 (0.404–0.423)<sup>1</sup> |
| **Total (cohort 2010)** |  |  |  |
| 1 | 12,792 | 517 | 0.960 (0.957–0.964)<sup>1</sup> |
| 2 | 12,275 | 873 | 0.892 (0.887–0.898)<sup>1</sup> |
| 3 | 11,402 | 908 | 0.820 (0.814–0.826)<sup>1</sup> |
| 4 | 10,494 | 812 | 0.756 (0.749–0.763)<sup>1</sup> |
| 5 | 9,682 | 702 | 0.701 (0.694–0.709)<sup>1</sup> |
| 6 | 8,980 | 684 | 0.647 (0.639–0.655)<sup>1</sup> |
| 7 | 8,296 | 648 | 0.593 (0.585–0.601)<sup>1</sup> |
| 8 | 7,648 | 647 | 0.544 (0.536–0.553)<sup>1</sup> |
| 9 | 7,001 | 649 | 0.495 (0.486–0.504)<sup>1</sup> |

**Supplementary Table 3:** 2000 and 2010 cohorts of scientists, IMMU, immunology and microbiology. Kaplan–Meier estimate by gender with total counts for men and women, time (in years), number of observations of scientists leaving science. Kaplan–Meier probability of staying in science with a 95% confidence interval. Note: (1) standard error of 0.01, (2) standard error of 0.05.

| Time (years) | Women (cohort 2000) |  |  | Men (cohort 2000) |  |  | Total (cohort 2000) |  |  | Probability of staying is higher for men by ... (%) |
| --- | --- | --- | --- | --- | --- | --- | --- | --- | --- | --- |
|  | n | n leaving science | KM probability (staying) with 95% CI and SE | n | n leaving science | KM probability (staying) with 95% CI and SE | n | n leaving science | KM probability (staying) with 95% CI and SE |  |
| 1 | 970 | 55 | 0.943 (0.929-0.958)<sup>1</sup> | 805 | 38 | 0.953 (0.953-0.953)<sup>1</sup> | 1,775 | 93 | 0.947 (0.939–0.955)<sup>1</sup> | 1.06 |
| 2 | 915 | 90 | 0.851 (0.828-0.873)<sup>2</sup> | 767 | 65 | 0.872 (0.872-0.872)<sup>2</sup> | 1,682 | 155 | 0.863 (0.852–0.874)<sup>2</sup> | 2.47 |
| 3 | 825 | 95 | 0.753 (0.726-0.780)<sup>2</sup> | 702 | 69 | 0.786 (0.786-0.786)<sup>2</sup> | 1,527 | 164 | 0.777 (0.764–0.789)<sup>2</sup> | 4.38 |
| 4 | 730 | 80 | 0.670 (0.641-0.700)<sup>2</sup> | 633 | 59 | 0.713 (0.713-0.713)<sup>2</sup> | 1,363 | 139 | 0.702 (0.688–0.716)<sup>2</sup> | 6.42 |
| 5 | 650 | 80 | 0.588 (0.557-0.619)<sup>2</sup> | 574 | 52 | 0.648 (0.648-0.648)<sup>2</sup> | 1,224 | 132 | 0.622 (0.608–0.637)<sup>2</sup> | 10.20 |
| 6 | 570 | 49 | 0.537 (0.507-0.569)<sup>2</sup> | 522 | 41 | 0.598 (0.598-0.598)<sup>2</sup> | 1,092 | 90 | 0.569 (0.555–0.584)<sup>2</sup> | 11.36 |
| 7 | 521 | 49 | 0.487 (0.456-0.519)<sup>2</sup> | 481 | 42 | 0.545 (0.545-0.545)<sup>2</sup> | 1,002 | 91 | 0.517 (0.502–0.533)<sup>2</sup> | 11.91 |
| 8 | 472 | 44 | 0.441 (0.411-0.474)<sup>2</sup> | 439 | 36 | 0.501 (0.501-0.501)<sup>2</sup> | 911 | 80 | 0.467 (0.451–0.483)<sup>2</sup> | 13.61 |
| 9 | 428 | 33 | 0.407 (0.377-0.439)<sup>2</sup> | 403 | 36 | 0.456 (0.456-0.456)<sup>2</sup> | 831 | 69 | 0.428 (0.412–0.444)<sup>2</sup> | 12.04 |
| 10 | 395 | 28 | 0.378 (0.349-0.410)<sup>2</sup> | 367 | 19 | 0.432 (0.432-0.432)<sup>2</sup> | 762 | 47 | 0.404 (0.388–0.420)<sup>2</sup> | 14.29 |
| 11 | 367 | 26 | 0.352 (0.323-0.383)<sup>2</sup> | 348 | 23 | 0.404 (0.404-0.404)<sup>2</sup> | 715 | 49 | 0.375 (0.359–0.392)<sup>2</sup> | 14.77 |
| 12 | 341 | 18 | 0.333 (0.305-0.364)<sup>2</sup> | 325 | 19 | 0.380 (0.380-0.380)<sup>2</sup> | 666 | 37 | 0.357 (0.340–0.374)<sup>2</sup> | 14.11 |
| 13 | 323 | 19 | 0.313 (0.286-0.344)<sup>2</sup> | 306 | 16 | 0.360 (0.360-0.360)<sup>2</sup> | 629 | 35 | 0.339 (0.322–0.356)<sup>2</sup> | 15.02 |
| 14 | 304 | 21 | 0.292 (0.265-0.322)<sup>2</sup> | 290 | 10 | 0.348 (0.348-0.348)<sup>2</sup> | 594 | 31 | 0.319 (0.303–0.336)<sup>2</sup> | 19.18 |
| 15 | 283 | 15 | 0.276 (0.250-0.306)<sup>2</sup> | 280 | 17 | 0.327 (0.327-0.327)<sup>2</sup> | 563 | 32 | 0.303 (0.287–0.320)<sup>2</sup> | 18.48 |
| 16 | 268 | 17 | 0.259 (0.233-0.288)<sup>2</sup> | 263 | 13 | 0.311 (0.311-0.311)<sup>2</sup> | 531 | 30 | 0.285 (0.269–0.302)<sup>2</sup> | 20.08 |
| 17 | 251 | 12 | 0.246 (0.221-0.275)<sup>2</sup> | 250 | 15 | 0.292 (0.292-0.292)<sup>2</sup> | 501 | 27 | 0.269 (0.253–0.286)<sup>2</sup> | 18.70 |
| 18 | 239 | 13 | 0.233 (0.208-0.261)<sup>2</sup> | 235 | 13 | 0.276 (0.276-0.276)<sup>2</sup> | 474 | 26 | 0.252 (0.236–0.268)<sup>2</sup> | 18.45 |
| 19 | 226 | 20 | 0.212 (0.188-0.240)<sup>2</sup> | 222 | 17 | 0.255 (0.255-0.255)<sup>2</sup> | 448 | 37 | 0.232 (0.216–0.249)<sup>2</sup> | 20.28 |
| Time (years) | Women (cohort 2010) |  |  | Men (cohort 2010) |  |  | Total (cohort 2010) |  |  | Probability of staying is higher for men by ... (%) |
|  | n | n leaving science | KM probability (staying) with 95% CI and SE | n | n leaving science | KM probability (staying) with 95% CI and SE | n | n leaving science | KM probability (staying) with 95% CI and SE |  |
| 1 | 1,422 | 68 | 0.952 (0.941-0.963)<sup>1</sup> | 939 | 61 | 0.935 (0.935-0.935)<sup>1</sup> | 2,361 | 129 | 0.945 (0.937–0.953)<sup>1</sup> | -1.79 |
| 2 | 1,354 | 151 | 0.846 (0.827-0.865)<sup>1</sup> | 878 | 90 | 0.839 (0.839-0.839)<sup>2</sup> | 2,232 | 241 | 0.843 (0.831–0.855)<sup>2</sup> | -0.83 |
| 3 | 1,203 | 140 | 0.748 (0.725-0.770)<sup>2</sup> | 788 | 73 | 0.761 (0.761-0.761)<sup>2</sup> | 1,991 | 213 | 0.753 (0.739–0.766)<sup>2</sup> | 1.74 |
| 4 | 1,063 | 142 | 0.648 (0.623-0.673)<sup>2</sup> | 715 | 80 | 0.676 (0.676-0.676)<sup>2</sup> | 1,778 | 222 | 0.662 (0.647–0.677)<sup>2</sup> | 4.32 |
| 5 | 921 | 107 | 0.572 (0.547-0.599)<sup>2</sup> | 635 | 62 | 0.610 (0.610-0.610)<sup>2</sup> | 1,556 | 169 | 0.590 (0.574–0.606)<sup>2</sup> | 6.64 |
| 6 | 814 | 98 | 0.504 (0.478-0.530)<sup>2</sup> | 573 | 53 | 0.554 (0.554-0.554)<sup>2</sup> | 1,387 | 151 | 0.528 (0.512–0.544)<sup>2</sup> | 9.92 |
| 7 | 716 | 91 | 0.440 (0.414-0.466)<sup>2</sup> | 520 | 57 | 0.493 (0.493-0.493)<sup>2</sup> | 1,236 | 148 | 0.468 (0.452–0.484)<sup>2</sup> | 12.05 |
| 8 | 625 | 73 | 0.388 (0.364-0.414)<sup>2</sup> | 463 | 50 | 0.440 (0.440-0.440)<sup>2</sup> | 1,088 | 123 | 0.415 (0.399–0.431)<sup>2</sup> | 13.40 |
| 9 | 552 | 77 | 0.334 (0.310-0.359)<sup>2</sup> | 413 | 51 | 0.386 (0.386-0.386)<sup>2</sup> | 965 | 128 | 0.363 (0.347–0.380)<sup>2</sup> | 15.57 |

Note: (1) standard error of 0.01, (2) standard error of 0.05.

**Supplementary Table 4:** 2010 and 2010 cohorts of scientists, NEURO, neuroscience. Kaplan–Meier estimate by gender with total counts for men and women, time (in years), number of observations of scientists leaving science. Kaplan–Meier probability of staying in science with a 95% confidence interval. Note: (1) standard error of 0.01, (2) standard error of 0.05.

| Time (years) | Women (cohort 2000) |  |  | Men (cohort 2000) |  |  | Total (cohort 2000) |  |  | Probability of staying is higher for men by ... (%) |
| --- | --- | --- | --- | --- | --- | --- | --- | --- | --- | --- |
|  | n | n leaving science | KM probability (staying) with 95% CI and SE | n | n leaving science | KM probability (staying) with 95% CI and SE | n | n leaving science | KM probability (staying) with 95% CI and SE |  |
| 1 | 1,180 | 67 | 0.943 (0.930-0.957)<sup>1</sup> | 1,353 | 41 | 0.970 (0.970-0.970)<sup>1</sup> | 2,533 | 108 | 0.957 (0.947–0.967)<sup>1</sup> | 2.86 |
| 2 | 1,113 | 97 | 0.861 (0.842-0.881)<sup>2</sup> | 1,312 | 97 | 0.898 (0.898-0.898)<sup>1</sup> | 2,425 | 194 | 0.880 (0.867–0.893)<sup>2</sup> | 4.30 |
| 3 | 1,016 | 87 | 0.787 (0.764-0.811)<sup>2</sup> | 1,215 | 93 | 0.829 (0.829-0.829)<sup>2</sup> | 2,231 | 180 | 0.805 (0.791–0.820)<sup>2</sup> | 5.34 |
| 4 | 929 | 95 | 0.707 (0.681-0.733)<sup>2</sup> | 1,122 | 74 | 0.775 (0.775-0.775)<sup>2</sup> | 2,051 | 169 | 0.740 (0.725–0.755)<sup>2</sup> | 9.62 |
| 5 | 834 | 86 | 0.634 (0.607-0.662)<sup>2</sup> | 1,048 | 55 | 0.734 (0.734-0.734)<sup>2</sup> | 1,882 | 141 | 0.682 (0.667–0.698)<sup>2</sup> | 15.77 |
| 6 | 748 | 61 | 0.582 (0.555-0.611)<sup>2</sup> | 993 | 59 | 0.690 (0.690-0.690)<sup>2</sup> | 1,741 | 120 | 0.632 (0.616–0.649)<sup>2</sup> | 18.56 |
| 7 | 687 | 34 | 0.553 (0.526-0.582)<sup>2</sup> | 934 | 47 | 0.656 (0.656-0.656)<sup>2</sup> | 1,621 | 81 | 0.598 (0.581–0.615)<sup>2</sup> | 18.63 |
| 8 | 653 | 47 | 0.514 (0.486-0.543)<sup>2</sup> | 887 | 49 | 0.619 (0.619-0.619)<sup>2</sup> | 1,540 | 96 | 0.561 (0.544–0.578)<sup>2</sup> | 20.43 |
| 9 | 606 | 42 | 0.478 (0.450-0.507)<sup>2</sup> | 838 | 42 | 0.588 (0.588-0.588)<sup>2</sup> | 1,444 | 84 | 0.529 (0.511–0.546)<sup>2</sup> | 23.01 |
| 10 | 564 | 34 | 0.449 (0.422-0.478)<sup>2</sup> | 796 | 33 | 0.564 (0.564-0.564)<sup>2</sup> | 1,360 | 67 | 0.504 (0.486–0.522)<sup>2</sup> | 25.61 |
| 11 | 530 | 23 | 0.430 (0.402-0.459)<sup>2</sup> | 763 | 40 | 0.534 (0.534-0.534)<sup>2</sup> | 1,293 | 63 | 0.480 (0.462–0.498)<sup>2</sup> | 24.19 |
| 12 | 507 | 29 | 0.405 (0.378-0.434)<sup>2</sup> | 723 | 29 | 0.513 (0.513-0.513)<sup>2</sup> | 1,230 | 58 | 0.458 (0.440–0.476)<sup>2</sup> | 26.67 |
| 13 | 478 | 28 | 0.381 (0.355-0.410)<sup>2</sup> | 694 | 31 | 0.490 (0.490-0.490)<sup>2</sup> | 1,172 | 59 | 0.437 (0.419–0.455)<sup>2</sup> | 28.61 |
| 14 | 450 | 17 | 0.367 (0.340-0.396)<sup>2</sup> | 663 | 19 | 0.476 (0.476-0.476)<sup>2</sup> | 1,113 | 36 | 0.423 (0.405–0.442)<sup>2</sup> | 29.70 |
| 15 | 433 | 24 | 0.347 (0.320-0.375)<sup>2</sup> | 644 | 24 | 0.458 (0.458-0.458)<sup>2</sup> | 1,077 | 48 | 0.403 (0.384–0.422)<sup>2</sup> | 31.99 |
| 16 | 409 | 20 | 0.330 (0.304-0.358)<sup>2</sup> | 620 | 22 | 0.442 (0.442-0.442)<sup>2</sup> | 1,029 | 42 | 0.386 (0.367–0.405)<sup>2</sup> | 33.94 |
| 17 | 389 | 27 | 0.307 (0.282-0.334)<sup>2</sup> | 598 | 22 | 0.426 (0.426-0.426)<sup>2</sup> | 987 | 49 | 0.365 (0.346–0.384)<sup>2</sup> | 38.76 |
| 18 | 362 | 24 | 0.286 (0.262-0.313)<sup>2</sup> | 576 | 26 | 0.407 (0.407-0.407)<sup>2</sup> | 938 | 50 | 0.345 (0.326–0.364)<sup>2</sup> | 42.31 |
| 19 | 338 | 15 | 0.274 (0.249-0.300)<sup>2</sup> | 550 | 33 | 0.382 (0.382-0.382)<sup>2</sup> | 888 | 48 | 0.326 (0.308–0.345)<sup>2</sup> | 39.42 |
| Time (years) | Women (cohort 2010) |  |  | Men (cohort 2010) |  |  | Total (cohort 2010) |  |  | Probability of staying is higher for men by ... (%) |
|  | n | n leaving science | KM probability (staying) with 95% CI and SE | n | n leaving science | KM probability (staying) with 95% CI and SE | n | n leaving science | KM probability (staying) with 95% CI and SE |  |
| 1 | 2,360 | 81 | 0.966 (0.958-0.973)<sup>1</sup> | 2,153 | 75 | 0.965 (0.965-0.965)<sup>1</sup> | 4,513 | 156 | 0.966 (0.961–0.971)<sup>1</sup> | -0.10 |
| 2 | 2,279 | 218 | 0.873 (0.860-0.887)<sup>1</sup> | 2,078 | 155 | 0.893 (0.893-0.893)<sup>1</sup> | 4,357 | 373 | 0.883 (0.873–0.894)<sup>1</sup> | 2.29 |
| 3 | 2,061 | 197 | 0.790 (0.774-0.806)<sup>1</sup> | 1,923 | 158 | 0.820 (0.820-0.820)<sup>1</sup> | 3,984 | 355 | 0.804 (0.792–0.817)<sup>1</sup> | 3.80 |
| 4 | 1,864 | 200 | 0.705 (0.687-0.724)<sup>1</sup> | 1,765 | 135 | 0.757 (0.757-0.757)<sup>1</sup> | 3,629 | 335 | 0.731 (0.718–0.745)<sup>1</sup> | 7.38 |
| 5 | 1,664 | 169 | 0.633 (0.614-0.653)<sup>2</sup> | 1,630 | 130 | 0.697 (0.697-0.697)<sup>1</sup> | 3,294 | 299 | 0.664 (0.650–0.679)<sup>2</sup> | 10.11 |
| 6 | 1,495 | 155 | 0.568 (0.548-0.588)<sup>2</sup> | 1,500 | 125 | 0.639 (0.639-0.639)<sup>2</sup> | 2,995 | 280 | 0.603 (0.588–0.618)<sup>2</sup> | 12.50 |
| 7 | 1,340 | 163 | 0.499 (0.479-0.519)<sup>2</sup> | 1,375 | 104 | 0.590 (0.590-0.590)<sup>2</sup> | 2,715 | 267 | 0.543 (0.528–0.558)<sup>2</sup> | 18.24 |
| 8 | 1,177 | 133 | 0.442 (0.423-0.463)<sup>2</sup> | 1,271 | 120 | 0.535 (0.535-0.535)<sup>2</sup> | 2,448 | 253 | 0.489 (0.474–0.505)<sup>2</sup> | 21.04 |
| 9 | 1,044 | 122 | 0.391 (0.371-0.411)<sup>2</sup> | 1,151 | 98 | 0.489 (0.489-0.489)<sup>2</sup> | 2,195 | 220 | 0.439 (0.423–0.455)<sup>2</sup> | 25.06 |

Note: (1) standard error of 0.01, (2) standard error of 0.05.

**Supplementary Figure 1:**
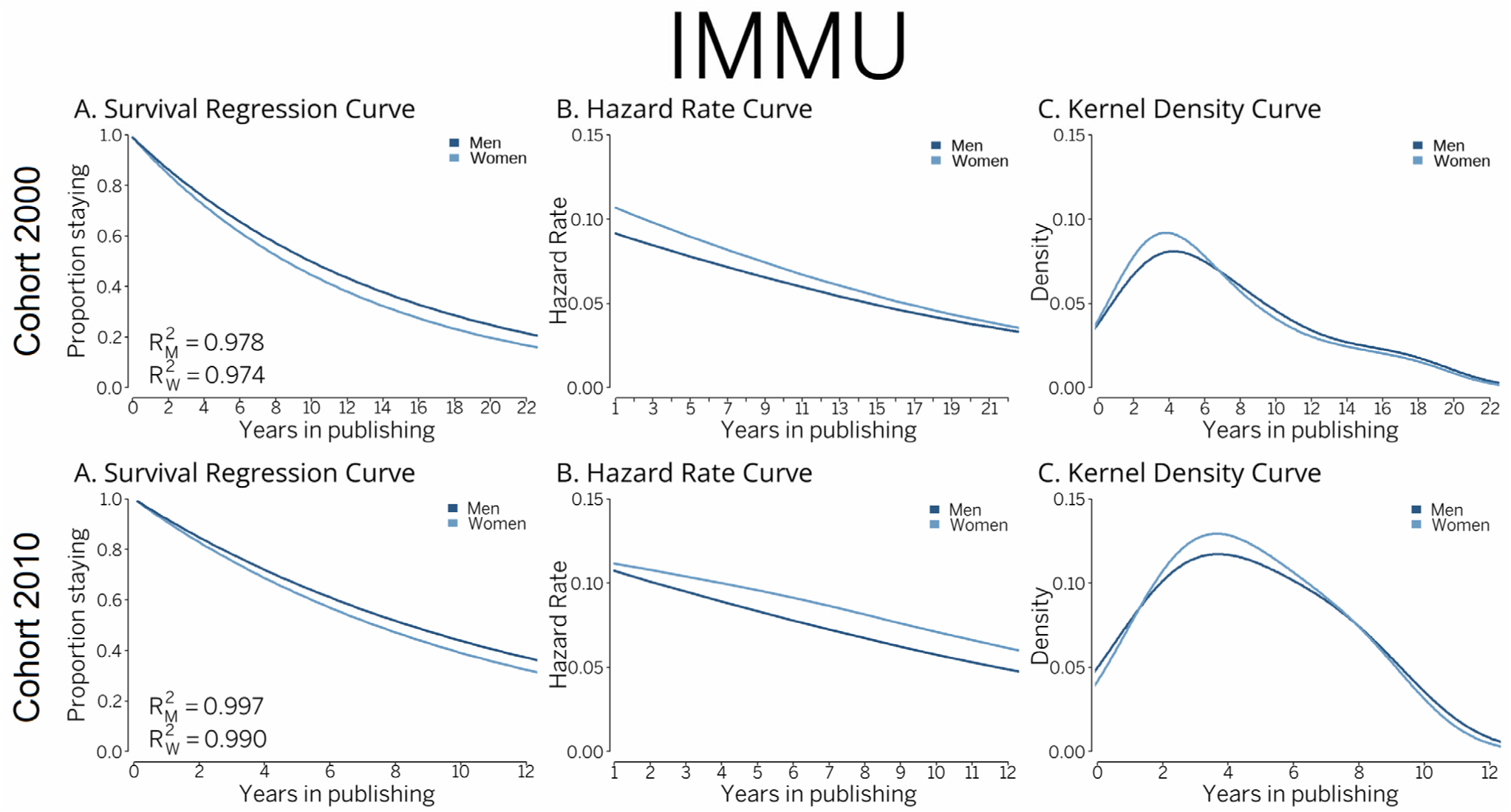
2000 (N=1,775) and 2010 (N=2,361) cohorts of scientists, IMMU, immunology and microbiology. Survival regression curve (exponential distribution fitting of Kaplan‒Meier curve), hazard rate curve (B-splines smoothing method, smoothing parameter 10k), and kernel density curve (B-splines smoothing method, bandwidth 2, component per point based on Gaussian curve)

**Supplementary Figure 2:**
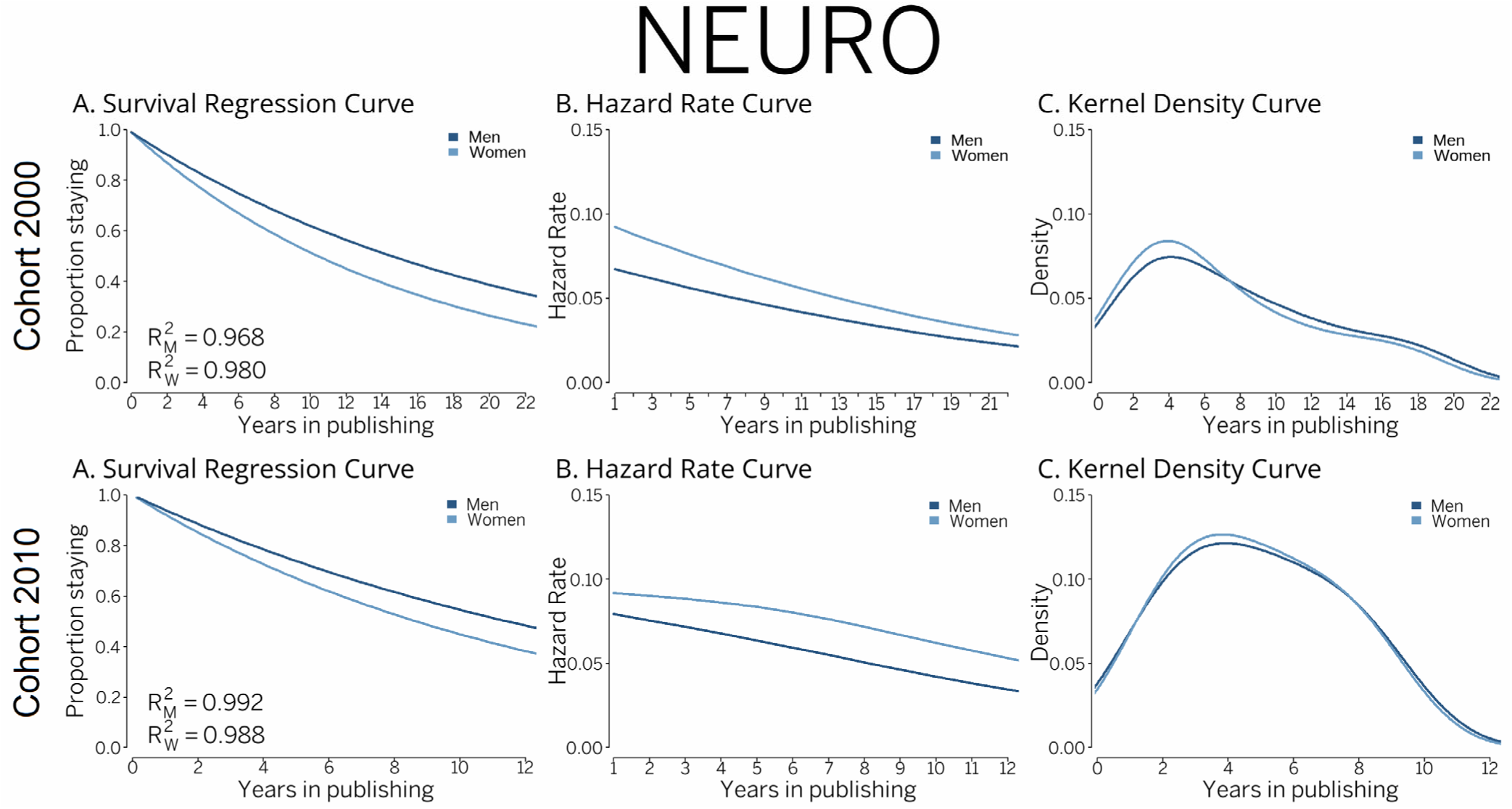
2000 (N=2,533) and 2010 (N=4,513) cohorts of scientists, NEURO, neuroscience. Survival regression curve (exponential distribution fitting of Kaplan‒Meier curve), hazard rate curve (B-splines smoothing method, smoothing parameter 10k), and kernel density curve (B-splines smoothing method, bandwidth 2, component per point based on Gaussian curve)

## References

1. Allison, P. D. (2014). Event history and survival analysis. Sage.

2. Baas, J., Schotten, M., Plume, A., Côté, G., & Karimi, R. (2020). Scopus as a curated, high-quality bibliometric data source for academic research in quantitative science studies. Quantitative Science Studies, 1(1), 377–386. 10.1162/qss_a_00019

3. Blickenstaff, J. C. (2005). Women and science careers: Leaky pipeline or gender filter? Gender and Education, 17(4), 369–386.

4. Branch, E. H. (Ed.) (2016). Pathways, potholes, and the persistence of women in science: Reconsidering the pipeline. Lexington Books.

5. Branch, E. H., & Alegria, S. (2016). Gendered responses to failure in undergraduate computing. Evidence, contradictions, and new directions. In E. H. Branch (Ed.), Pathways, potholes, and the persistence of women in science. Reconsidering the pipeline (pp. 17–31). Lexington Books.

6. Cornelius, R., Constantinople, A., & Gray, J. (1988). The chilly climate: Fact or artifact? The Journal of Higher Education, 59(5), 527–555.

7. Ehrenberg, R. G., Kasper, H., & Rees, D. I. (1991). Faculty turnover in American colleges and universities. Economics of Education Review, 10(2), 99–110.

8. Fox, M. F. (2010). Women and men faculty in academic science and engineering: Social-organizational indicators and implications. American Behavioral Scientist, 53, 997–1012.

9. Fox, M. F. (2020). Gender, science, and academic rank: Key issues and approaches. Quantitative Science Studies, 1(3), 1001–1006.

10. Fox, M. F., & Mohapatra, S. (2007). Social-organizational characteristics of work and publication productivity among academic scientists in doctoral-granting departments. The Journal of Higher Education, 78(5), 542–571.

11. Fox, M. F., & Kline, K. (2016). Women faculty in computing. A key case of women in science. In E. H. Branch (Ed.), Pathways, potholes, and the persistence of women in science: Reconsidering the pipeline (pp. 54–69). Lexington Books.

12. Geuna, A., & Shibayama, S. (2015). Moving out of academic research: Why do scientists stop doing research? In A. Geuna (Ed.), Global mobility of research scientists (pp. 271–297). Elsevier.

13. Glenn, N. D. (2005). Cohort analysis. Sage.

14. Hermanowicz, J. (2012). The sociology of academic careers: Problems and prospects. In J. C. Smart & M. B. Paulsen (Eds.), Higher education: Handbook of theory and research (Vol. 27). Springer.

15. Kaminski, D., & Geisler, C. (2012). Survival analysis of faculty retention in science and engineering by gender. Science, 335, 864–866.

16. Karimi, F., Wagner, C., Lemmerich, F., Jadidi, M., & Strohmaier, M. (2016). Inferring gender from names on the web: A comparative evaluation of gender detection methods. In Proceedings of the 25th International Conference Companion on World Wide Web (pp. 53–54).

17. Kashyap, R., Rinderknecht, R. G., Akbaritabar, A., Alburez-Gutierrez, D., Gil-Clavel, S., Grow, A., & Zhao, X. (2023). Digital and computational demography. In Research handbook on digital sociology (pp. 48–86). Edward Elgar Publishing.

18. Kwiek, M. (2019). Changing European academics: A comparative study of social stratification, work patterns and research productivity. Routledge.

19. Kwiek, M., & Roszka, W. (2021a). Gender disparities in international research collaboration: A large-scale bibliometric study of 25,000 university professors. Journal of Economic Surveys, 35(5), 1344–1388.

20. Kwiek, M., & Roszka, W. (2021b). Gender-based homophily in research: A large-scale study of man-woman collaboration. Journal of Informetrics, 15(3), 1–38.

21. Kwiek, M., & Roszka, W. (2022). Are female scientists less inclined to publish alone? The gender solo research gap. Scientometrics, 127, 1697–1735.

22. Kwiek, M., & Roszka, W. (2024). Are scientists changing their research productivity classes when they move up the academic ladder? Innovative Higher Education, 1–39. Online first: 10.1007/s10755-024-09735-3

23. Kwiek, M., & Szymula, L. (2023). Young male and female scientists: A quantitative exploratory study of the changing demographics of the global scientific workforce. Quantitative Science Studies, 4(4), 902–937.

24. Kwiek, M., & Szymula, L. (2024). Quantifying attrition in science: A cohort-based, longitudinal study of scientists in 38 OECD countries. Higher Education, 1–29. Online first: 10.1007/s10734-024-01284-0

25. Leisyte, L., & Dee, J. R. (2012). Understanding academic work in a changing institutional environment. In J. C. Smart & M. B. Paulsen (Eds.), Higher education: Handbook of theory and research (Vol. 27, pp. 123–206). Springer Netherlands.

26. Liu, L., Jones, B. F., Uzzi, B., et al. (2023). Data, measurement, and empirical methods in the science of science. Nature Human Behaviour, 7, 1046–1058.

27. Mills, M. (2011). Introducing survival and event history analysis. Sage.

28. Milojevic, S., Radicchi, F., & Walsh, J. P. (2018). Changing demographics of scientific careers: The rise of the temporary workforce. Proceedings of the National Academy of Sciences, 115, 12616–12623.

29. Naddaf, M. (2024, October 3). Nearly 50% of researchers quit science within a decade, huge study reveals. Nature News.

30. NamSor. (2024). NamSor API documentation. https://namsor.app/api-documentation/.

31. Preston, A. E. (2004). Leaving science: Occupational exit from scientific careers. Russell Sage Foundation.

32. Rosser, V. J. (2004). Faculty members’ intentions to leave: A national study on their work-life and satisfaction. Research in Higher Education, 45(3), 285–309.

33. Salganik, M. J. (2018). Bit by bit: Social research in the digital age. Princeton University Press.

34. Sebo, P. (2021). Performance of gender detection tools: A comparative study of name-to-gender inference services. Journal of the Medical Library Association, 109(3), 414–421.

35. Shaw, A. K., & Stanton, D. E. (2012). Leaks in the pipeline: Separating demographic inertia from ongoing gender differences in academia. *Proceedings of the Royal Society B*. Biological Sciences, 279(1743), 3736–3741.

36. Shibayama, S., & Baba, Y. (2015). Impact-oriented science policies and scientific publication practices: The case of life sciences in Japan. Research Policy, 44(4), 936–950.

37. Singer, J. D., & Willett, J. B. (2003). Applied longitudinal data analysis. Modeling change and event occurrence. Oxford University Press.

38. Smart, J. C. (1990). A causal model of faculty turnover intentions. Research in Higher Education, 31(5), 405–424.

39. Spoon, K., LaBerge, N., Wapman, K. H., Zhang, S., Morgan, A. C., Galesic, M., Fosdick, B. K., Larremore, D. B., & Clauset, A. (2023). Gender and retention patterns among U.S. faculty. Science Advances, 9, eadi2205. 10.1126/sciadv.adi2205

40. Sugimoto, C., & Larivière, V. (2023). Equity for women in science: Dismantling systemic barriers to advancement. Harvard University Press.

41. Wang, D., & Barabási, A. L. (2021). The science of science. Cambridge University Press.

42. White-Lewis, D. K., O’Meara, K., Mathews, K., et al. (2023). Leaving the institution or leaving the academy? Analyzing the factors that faculty weigh in actual departure decisions. Research in Higher Education, 64, 473–494.

43. Wolfinger, N. H., Mason, M. A., & Goulden, M. (2008). Problems in the pipeline: Gender, marriage, and fertility in the ivory tower. Journal of Higher Education, 79(4), 388–405.

44. Xie, Y., & Shauman, K. A. (2003). Women in science: Career processes and outcomes. Harvard University Press.

45. Zhou, Y., & Volkwein, J. F. (2004). Examining the influence on faculty departure intentions: A comparison of tenured versus nontenured faculty at research universities using NSOPF-99. Research in Higher Education, 45(2), 139–176.

